# Information maximization explains state-dependent synaptic plasticity and memory reorganization during non-rapid eye movement sleep

**DOI:** 10.1101/2022.03.22.485283

**Authors:** Kensuke Yoshida, Taro Toyoizumi

**Author notes:** Correspondence should be addressed to Kensuke Yoshida at or Taro Toyoizumi at.

## Abstract

Slow waves during the non-rapid eye movement (NREM) sleep reflect the alternating up and down states of cortical neurons; global and local slow waves promote memory consolidation and forgetting, respectively. Furthermore, distinct spike-timing-dependent plasticity (STDP) operates in these up and down states. The contribution of different plasticity rules to neural information coding and memory reorganization remains unknown. Here, we show that optimal synaptic plasticity for information maximization in a cortical neuron model provides a unified explanation for these phenomena. The model indicates that the optimal synaptic plasticity is biased towards depression as the baseline firing rate increases. This property explains the distinct STDP observed in the up and down states. Furthermore, it explains how global and local slow waves predominantly potentiate and depress synapses, respectively, if the background firing rate of excitatory neurons declines with the spatial scale of waves as the model predicts. The model provides a unifying account of the role of NREM sleep, bridging neural information coding, synaptic plasticity, and memory reorganization.

## 1 Introduction

Sleep is an essential physiological process and is widely conserved across species. One proposed role of sleep is to reorganize memory by regulating synaptic plasticity; some memories of awake experiences are consolidated, whereas others are forgotten [1, 2, 3]. Multiple studies have explored the mechanism behind memory consolidation and forgetting by focusing on slow waves observed during non-rapid eye movement (NREM) sleep. Slow waves are low-frequency (< 4 Hz) waves in electroencephalography (EEG) and local field potential (LFP), and their presence distinguishes NREM sleep from awake or rapid eye movement (REM) sleep. Each cortical neuron shows low-frequency transitions between the up (depolarized membrane potential) and down (hyperpolarized membrane potential) states synchronous to slow waves [4]. Both the correlational and causal relationships between slow waves and memory consolidation have been established. It has been reported that slow-wave activity is correlated with task performance after sleep [5], and boosting slow waves can enhance memory consolidation [6, 7, 8, 9].

Several studies have suggested that slow waves should be separated into distinct classes [10, 11, 12, 13, 14, 15, 16]. Although different classification schemes have been used in previous studies, one of the classes is more global, while the other is more local [10, 12, 13, 14]. A recent study further suggested that these two classes of slow waves have opposite effects on memory reorganization; the global and local classes promote memory consolidation and forgetting, respectively [13]. These studies suggest that memory reorganization is induced depending on the subtle sleep states such as the up and down states of global and local slow waves.

One possible explanation for how these sleep states differentially modulate the memory reorganization is that synaptic plasticity is modulated depending on the sleep state. Previous studies have shown that neuronal activity patterns in the awake state are reactivated within the slow waves during NREM sleep [17, 18, 19]. Although the synaptic plasticity rule during NREM sleep is largely unknown, a recent experimental study using anesthetized young mice *in vivo* has suggested that the spike-timing-dependent plasticity (STDP) during up states is biased toward depression compared with down states [20]. Consistently, another experimental study using acute brain slices demonstrated that the subthreshold inputs during up but not down states induce synaptic weakening [21]. These findings suggest that neuronal reactivation can induce different synaptic plasticity in the up and down states. This difference might be the key to understanding memory reorganization during NREM sleep and raises two further issues worth exploring theoretically. First, what is the benefit of modulating the synaptic plasticity rule depending on the up and down states? Because the nervous system has evolved to work efficiently, the efficiency of neuronal coding might be enhanced by this modulation. Second, how does the state-dependent synaptic plasticity reorganize memories in global and local slow waves?

To understand these issues, we adopted a normative approach based on the information maximization (infomax) principle [22, 23] and derived a synaptic plasticity rule for a spiking neuron model [24, 25] that achieves efficient information transmission. We found that the baseline firing rate is an important parameter of the infomax rule. An increased baseline firing rate biases the synaptic plasticity towards depression, consistent with the reported difference in STDP between the up and down states. We then constructed a neuronal network model exhibiting global and local slow waves and showed that four states (up and down states of global and local slow waves) have distinct STDP owing to different baseline firing rates. Finally, we suggest that the difference in synaptic plasticity in global and local slow waves can set a balance between memory consolidation and forgetting, consistent with the previous experimental findings.

## 2 Result

### 2.1 Optimal synaptic plasticity is biased toward depression in high firing rates

To consider optimal synaptic plasticity in different sleep states, we first considered a feedforward network model with a postsynaptic neuron and multiple excitatory presynaptic neurons. In this model, presynaptic spikes at synapse *j* evoked excitatory postsynaptic potentials (EPSPs) with the amplitude *w*_*j*_ and exponential decays with a time constant of 25 ms. The membrane potential of the postsynaptic neuron was computed as *u*(*t*) = *u*_*r*_ + Σ_*j*_ *w*_*j*_*h*_*j*_(*t*), where *u*_*r*_ is the resting membrane potential and *h*_*j*_ is the EPSP time-course from presynaptic neuron *j* with an instantaneous increment of 1 after each presynaptic spike. The postsynaptic neuron emits spikes with firing probability density *g*(*u*(*t*))*R*(*t*), where *g*(*u*) is a softplus activation intensity function, and refractory factor *R*(*t*) models the transient suppression of the postsynaptic firing rate after a postsynaptic spike (see Methods). Using the infomax approach, we assumed that the synaptic weights *w*_*j*_ change following the gradient of the mutual information between the presynaptic and postsynaptic spikes under the constraint of the synaptic weight cost [24, 25]. Thus, the synaptic weight changes were described by 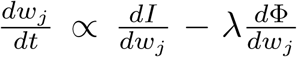 with mutual information *I* and synaptic weight cost Φ (see Methods). Coefficient *λ* controls the importance of the synaptic cost term relative to the information term. We omitted the homeostatic term assumed in the previous studies because it did not contribute to our results, where the postsynaptic firing rate was kept on average within the homeostatic range. The gradient of mutual information *dI/dw*_*j*_ was explicitly derived, and can be computed in real-time using only the variables observable at synapse *j*, namely, the EPSP time-course *h*_*j*_, postsynaptic spikes informed by a back-propagating action potential, postsynaptic activation intensity *g*(*u*) as a function of postsynaptic membrane potential *u*, refractory factor *R*, and mean activation intensity 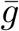 (see Methods). In addition, the gradient of the cost term decreased the synaptic strength by *λw*_*j*_ for every presynaptic spike of neuron *j*.

We ran simulations imitating the experimental STDP protocols *in vivo* [20]. To mimic the experimental setup, we divided the presynaptic neurons into the stimulated and non-stimulated neurons (Fig. 1A). Twenty stimulated neurons synchronously emitted a spike upon external presynaptic stimulation, and their synaptic weights changed according to the infomax rule. One hundred non-stimulated neurons spontaneously emitted Poisson spikes at 2.0 and 0.1 Hz in the up and down states, respectively, while their synaptic weights were fixed for simplicity. The mean activation intensity, 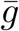, was computed by taking the average of *g*(*u*) in each state. This yielded 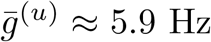 and 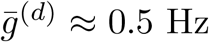 for the up and down states, respectively.

**Figure 1.**
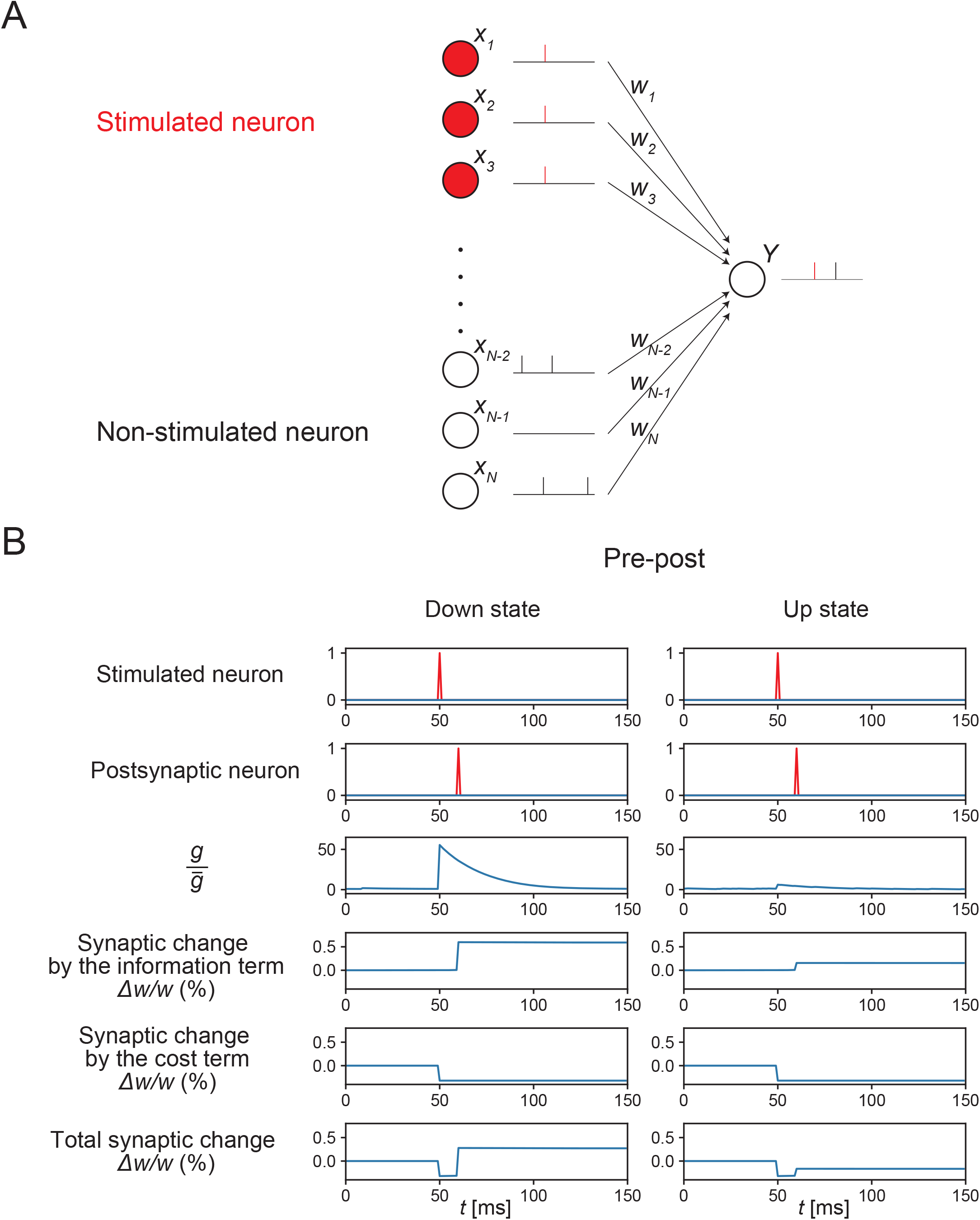
The infomax rule in the feedforward model. (A) A feedforward model of synaptic plasticity. Stimulated neurons synchronously emit spikes upon external presynaptic stimulation, and their synaptic weights change according to the infomax rule, whereas non-stimulated neurons spontaneously emit Poisson spikes and their synaptic weights are fixed for simplicity. A postsynaptic neuron emits both spontaneous spikes and evoked spikes upon external postsynaptic stimulation. The red vertical bars represent evoked spikes. (B) Representative traces of a stimulated neuron’s activity, the postsynaptic neuron’s activity, the ratio of the momentary and mean activation intensity 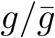, and changes of a synaptic weight from a stimulated neuron in the down and up states. Synaptic changes by the infomax rule were computed by summing the effects of the information term 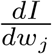 and cost term 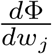. The synaptic increase by the information term was smaller in the up state than that in the down state.

We first characterized synaptic changes induced by pre-post stimulation, where a presynaptic spike was induced 10 ms before the postsynaptic spike. Representative traces of a synaptic weight from a stimulated neuron in the up and down states are plotted in Fig. 1B. The increase in 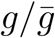 after presynaptic stimulation was greater when the mean firing rate was low, indicating that a postsynaptic spike can transmit a greater amount of information at a lower mean firing rate. Consequently, the information term caused a greater synaptic potentiation in the down state than that in the up state. The amount of synaptic potentiation due to the information term was roughly proportional to 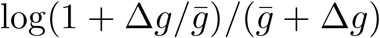, where 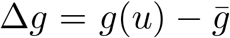 represents the increment in activation intensity due to the presynaptic stimulation (see Methods for details). Intuitively, Δ*g* measured the reliability of a synapse for transmitting the signal, and 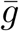 represented the noise level that quantifies the frequency of the postsynaptic spikes in the background. Thus, 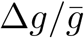 corresponds to the signal-to-noise ratio. By contrast, the change in the synaptic weight by the cost term was *λw*_*j*_ after every presynaptic spike of neuron *j*, regardless of the mean firing rate.

Figure 1B displays the representative traces of synaptic weight; however, synaptic changes also depended on other postsynaptic spikes and presynaptic spikes from non-stimulated neurons, which can occur randomly. Below, we quantify the average synaptic changes induced by three kinds of stimulation: pre-only stimulation and post-pre-stimulation (a presynaptic spike was induced 10 ms after an induced postsynaptic spike), in addition to the pre-post stimulation explored above. We started by simulating the pre-only stimulation. Experimentally, pre-only stimulation in the down state did not significantly change the synaptic weights of stimulated neurons [20]. To reproduce this experimental result, we set the coefficient of the cost term to *λ* = 0.32 (mV)^*−*2^, so that the changes in the stimulated synapses were on average zero in the down state (Fig. 2A). This value of *λ* was used throughout this paper. Simulations of the up state showed an overall synaptic depression because the synaptic potentiation due to the information term decreased with the mean firing rate for the reason described above.

**Figure 2.**
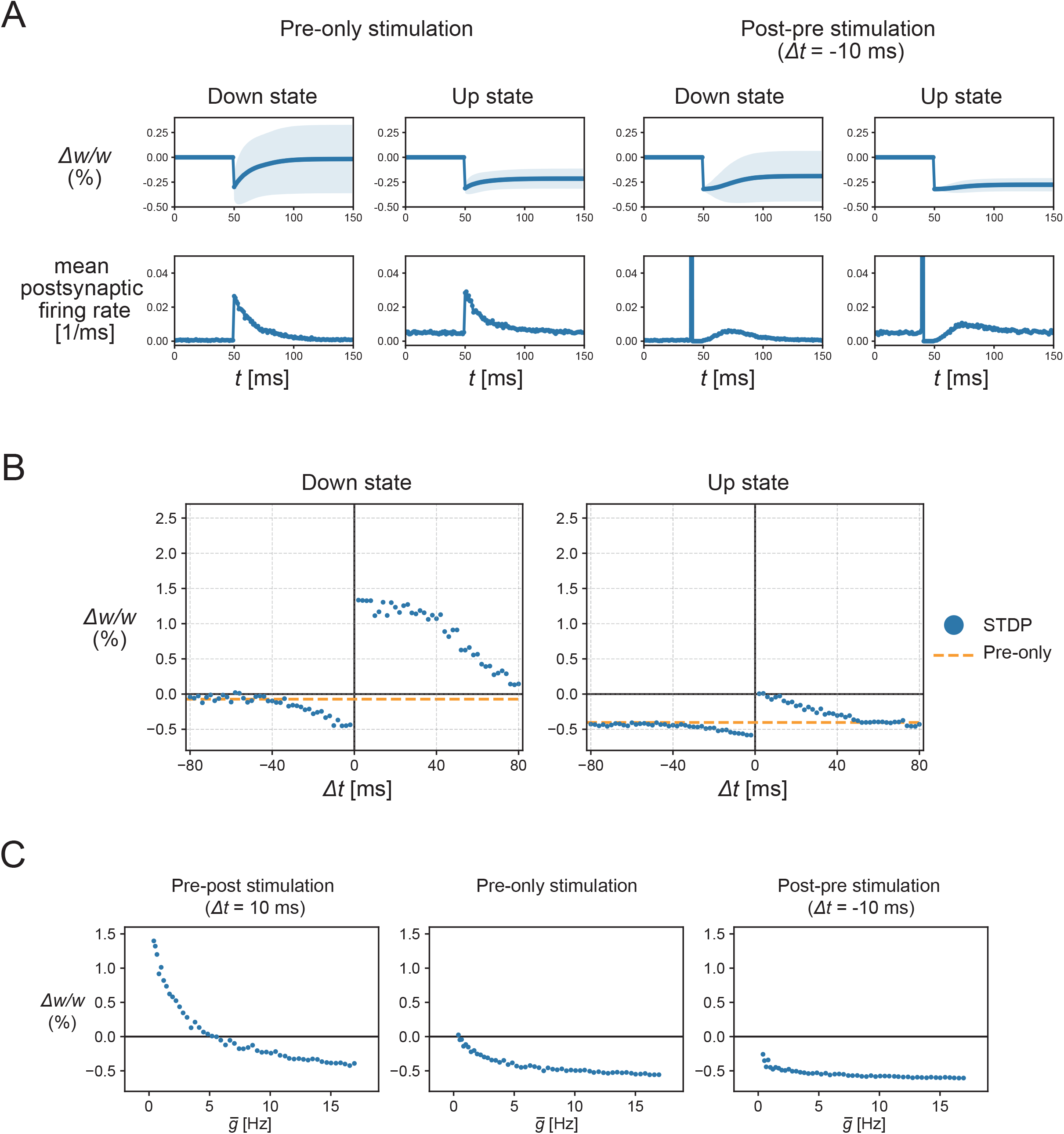
The synaptic plasticity induced by the STDP stimulations. (A) The mean traces of a synaptic weight and the mean postsynaptic firing rate in the pre-only stimulations or the post-pre-stimulations with Δ*t* = −10 ms, where the stimulated neurons emitted a synchronous spike upon the external stimulation at *t* = 50 ms. In the post-pre-stimulations, the post-synaptic neuron emitted an evoked spike at *t* = 40 ms and also responded to the presynaptic stimulation at *t* = 50 ms with some delay due to refractoriness. After the pre-only stimulations, the synaptic weight changed little in the down state, but was depressed in the up state. In the post-pre-stimulation, the synaptic weight was depressed in both the down and up states. The lines and shadows of the weight traces represent the means and standard deviations, respectively. (B) The synaptic changes by the STDP stimulations (blue points) and pre-only stimulations (orange dotted lines). Synaptic plasticity was biased towards depression in the up state. (C) Synaptic changes were dependent on mean activation intensity. The synaptic changes in pre-post stimulation with Δ*t* = 10 ms and pre-only stimulation decreased with increasing mean activation intensity, whereas the synaptic changes in post-pre-stimulation with Δ*t* = −10 ms were less insensitive to mean activation intensity.

Next, if a postsynaptic spike was evoked before the presynaptic stimulation (i.e., post-pre-stimulation), the infomax rule caused the synaptic depression both in the up and down states because the induced postsynaptic spike before the presynaptic stimulation suppressed *R*(*t*) and prevented the synaptic weights from increasing by the information term, whereas the cost term could still decrease these synapses (Fig. 2A).

To investigate how the STDP window of the infomax rule differs in the up and down states, the time difference between presynaptic and postsynaptic stimulations was systematically changed. Consistent with the observations above, the entire STDP curve was biased toward synaptic depression in the high mean firing rate condition (Fig. 2B). (See, also Fig. S2 for more systematic parameter dependency) While the synaptic change caused by post-pre-stimulation was relatively insensitive to mean firing rates, the high mean firing rates biased the synaptic changes toward depression with the pre-post and pre-only stimulations (Fig. 2C). These results were consistent with the corresponding *in vivo* experimental results [20]. Hence, the modulation of the infomax rule by the mean firing rate explained the synaptic plasticity during the up and down states in slow waves.

### 2.2 Firing rates of excitatory neurons are higher during local slow waves than those during global slow waves

To study how the above-mentioned findings would apply to memory reorganization during NREM sleep, we constructed the network models of cortical neurons that generate slow waves. We started by constructing a spatially homogeneous model similar to that in [26] and then introduced spatial heterogeneity to produce local and global slow waves. The network model consisted of recurrently connected spiking neurons, including the excitatory and inhibitory neurons (see Methods). The spatially homogeneous model assumed no connections between two inhibitory neurons but all-to-all connections between two excitatory neurons, and between excitatory and inhibitory neurons. Each excitatory neuron had adaptation currents that accumulated with the spikes. Adaptation currents correspond to, for example, the potassium currents involved in generating slow waves [27, 28, 29, 30]. The activation functions of excitatory and inhibitory neurons were both modeled by softplus functions, but the threshold and slope were greater for the inhibitory neurons than those for the excitatory neurons (Fig. 3B). In these settings, the excitatory and inhibitory activities showed bistability of the up and down states, and the transitions were caused by the slowly changing adaptation currents (Fig. S1A-B). This was consistent with the previous theoretical study [26]. The inhibitory population was mostly inactive in down states because of the high threshold (Fig. 3B), but was active in the up state and stabilized the recurrent excitatory activity.

**Figure 3.**
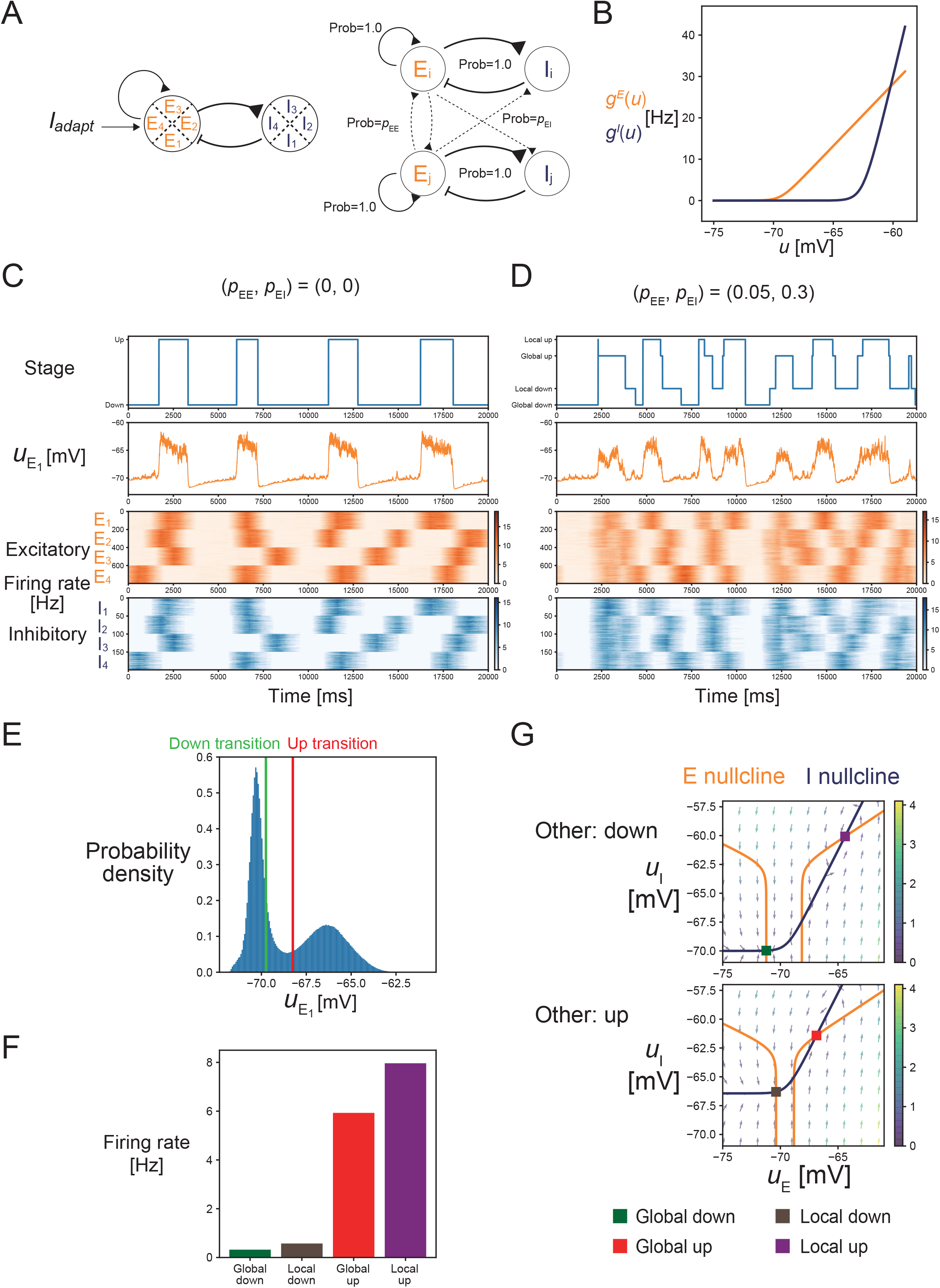
The cortical network model exhibiting global and local slow waves (A) The schematic description of the model. (Left) Model composed of four local networks. Each local network included 200 excitatory and 50 inhibitory neurons. (Right) Within a network, connections existed between two excitatory neurons, and between excitatory and inhibitory neurons, while two inhibitory neurons had no connections between them. Between different networks, there were sparse connections from excitatory to excitatory or inhibitory neurons but none from the inhibitory neurons. (B) The activation functions of excitatory and inhibitory neurons. Both functions were softplus functions, but the threshold and slope were greater for inhibitory neurons than those for the excitatory neurons. (C) The dynamics of the slow wave model in the case of no connections between different local networks. The up- and down-states of the E1 population were classified using the mean membrane potential 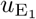. The four networks independently transited between the up and down states. (D) The dynamics of the slow wave model in the case that there exist sparse connections between different networks. The up and down states of the E1 population were classified into the global or local states depending on the states of the other populations. (E) The probability density of the membrane potential. The transition thresholds to the up and down states are indicated by red and green lines, respectively. (F) The mean firing rates of the E1 population in global down, local down, global up, and local up states. (G) The phase plane in the case that other populations are in down states or in up states. Excitatory and inhibitory nullclines are shown as orange and blue lines, respectively.

To consider the difference between global and local slow waves, we extended the model by embedding four local networks, each as described above, within the overall network (Fig. 3A). In cases of no between-network connections, each local network independently produced up and down cycles of slow waves (Fig. 3C). Next, we introduced sparse long-range excitatory connections between different local networks. We assumed that the long-range connections project to both the excitatory and inhibitory neurons, as demonstrated in previous modeling studies [31]. In this setting, each local network showed the transitions between the up and down states, some of which were local, whereas others were global in synchrony across the local networks (Fig. 3D). To objectively define global and local slow waves, we first classified the up and down states of each local network based on the mean membrane potential averaged across excitatory neurons (Fig. 3C, E). Transitions to the down and up states were detected when the mean membrane potential decreased below -69.75 mV and exceeded -68.25 mV, respectively. We classified each state into a global or local state by counting the number of up and down states across the local networks (Fig. 3D) (see Methods). In the global up/down states, either all or all but one local network simultaneously achieved the same state.

We then analyzed how the difference between global and local slow waves affected the learning by the infomax rule. Since the outcome of the infomax rule depends on the mean firing rates, we examined the mean firing rates in the up and down states during the global and local slow waves. The mean firing rates of excitatory neurons followed global down < local down < global up < local up in ascending order in the simulations (Fig. 3F). The difference between the global and local down states was simply explained by the strength of the long-range excitation from the surrounding networks to the local excitatory population. Because the surrounding excitatory populations had elevated activity in their up state, the long-range excitation was stronger in the local down states than that in the global down states. Note that the local inhibitory population was mostly inactive in both the local and global down states, and did not contribute significantly to the difference. By contrast, the difference between the global and local up states was mainly explained by the strength of the local inhibition to the excitatory population. While the local network was in the up state, its inhibitory activity was more sensitive to the long-range excitation than its excitatory activity because of the steeper inhibitory activation function at high membrane potential (see Fig. 3B and Methods). Therefore, the strong long-range excitation from the surrounding networks in the global up state effectively reduced the local excitatory activity via elevated local inhibition. To verify this, we performed a phase plane analysis, assuming a large number of neurons (see Methods). The phase planes showed that the firing rates of the two stable points (i.e., up and down states) changed depending on the state of the surrounding networks (Fig. 3G). As expected, the membrane potential of the excitatory neurons was higher in local down states with elevated long-range excitation than that in the global down states. In addition, the membrane potential of excitatory neurons was lower in the global up states with elevated local inhibition than that in the local up states. In this case, the long-range excitation shifted both the excitatory and inhibitory nullclines. Since the shift of the inhibitory nullcline was much larger than that of the excitatory nullcline, the membrane potential of excitatory neurons decreased with the long-range excitation (Fig. 3G). The observed higher local excitatory activity in the local up states than that of the global up states is a natural consequence of unstable recurrent excitatory dynamics being stabilized by the local inhibitory population. A similar response to external input was previously demonstrated both experimentally and theoretically as a property of inhibition-stabilized networks (ISNs) [32, 33, 31]. Based on these results, we hypothesized that the infomax rule, which is sensitive to the baseline firing rate, would yield distinct learning outcomes in the global and local slow waves.

### 2.3 Optimal synaptic plasticity in up and down states of global and local slow waves

To study the outcome of the infomax rule in global and local slow waves, we first explored the STDP window using the slow-wave model in the previous section. We introduced 20 presynaptic excitatory neurons that spike synchronously when externally stimulated, as shown in Fig. 2. These presynaptic neurons projected to a randomly selected postsynaptic neuron in the first excitatory population, E1, of the slow-wave model (Fig. 4A). We assumed that these feedforward synaptic weights were updated by the infomax rule, whereas the recurrent synaptic weights were fixed. The changes in the feedforward synaptic weights averaged over random realizations of the model are shown below.

**Figure 4.**
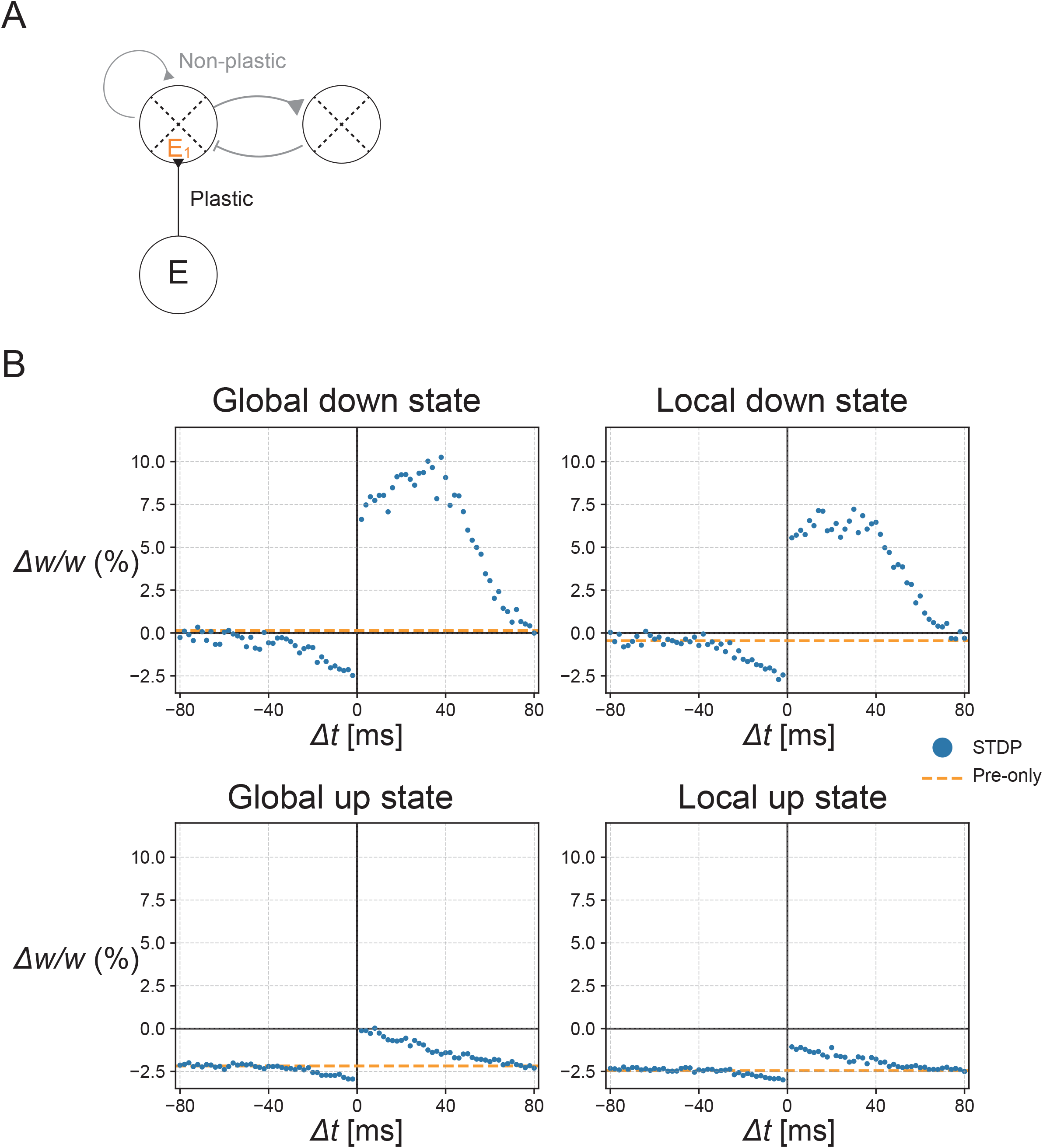
STDP in different sleep states. (A) The schematic description of the simulation. Presynaptic neurons had a feedforward projection onto an excitatory postsynaptic neuron in the E1 population. The feedforward synaptic weights were plastic, whereas the recurrent synaptic weights were fixed. (B) The synaptic changes caused by the STDP stimulations (blue points) and pre-only stimulations (orange dotted lines) in each sleep state. The amount of synaptic changes follows global down *>* local down *>* global up *>* local up, in descending order.

We first studied the STDP by evoking a postsynaptic spike at a fixed time, before or after the presynaptic stimulation. In addition to this evoked spike, the postsynaptic neuron could generate other spikes triggered by the network activity. In this model, the mean activation intensity 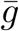 was computed by averaging the activation intensity *g*(*u*_*i*_) of neuron *i* for neurons included within the same excitatory population (see Discussion for possible biological implementations). As expected from Figs. 2 and 3, the STDP results depended on the sleep states of the slow-wave model (Fig. 4B). These results indicate two important points. First, the synaptic plasticity in the up states is biased toward synaptic depression as compared with the down states, which is consistent with the experimental findings [20]. Second, synaptic plasticity in the local up and down states is biased toward synaptic depression as compared with the global up and down states, respectively. This property results from the model prediction that the mean firing rates are higher in the local up and down states than those in the corresponding global states.

To further investigate the impact of sleep states on memory reorganization, we simulate how synaptic weights that contribute to task performance change during subsequent sleep. This time, we simulated 40 presynaptic neurons that emitted spikes synchronously upon the presentation of a task cue and projected to a postsynaptic neuron in the E1 population of the slow-wave model (Fig. 5A). The simulation was repeated over random realizations of the model parameters. Inspired by the brain-machine-interface task [13], we defined task performance as an increase in the postsynaptic firing rate upon the presentation of a task cue. We considered the synaptic changes during post-learning NREM sleep, assuming that the feedforward synaptic weights have already been potentiated to elevate the postsynaptic firing rate during the task. We updated the feedforward synaptic weights during post-learning NREM sleep according to the infomax rule. Presynaptic neurons synchronously emit spikes with Poisson statistics of 5 Hz when the memory of the task is reactivated [17, 19]. We restricted memory reactivation to occur during the up states of the local network (Fig. 5B) for easy comparison with the experimental results by Kim et al. [13], which focused on the role of reactivation in up states. (Note that memory reactivation might also exist in down states [34].) Further simulation details are described in Methods.

**Figure 5.**
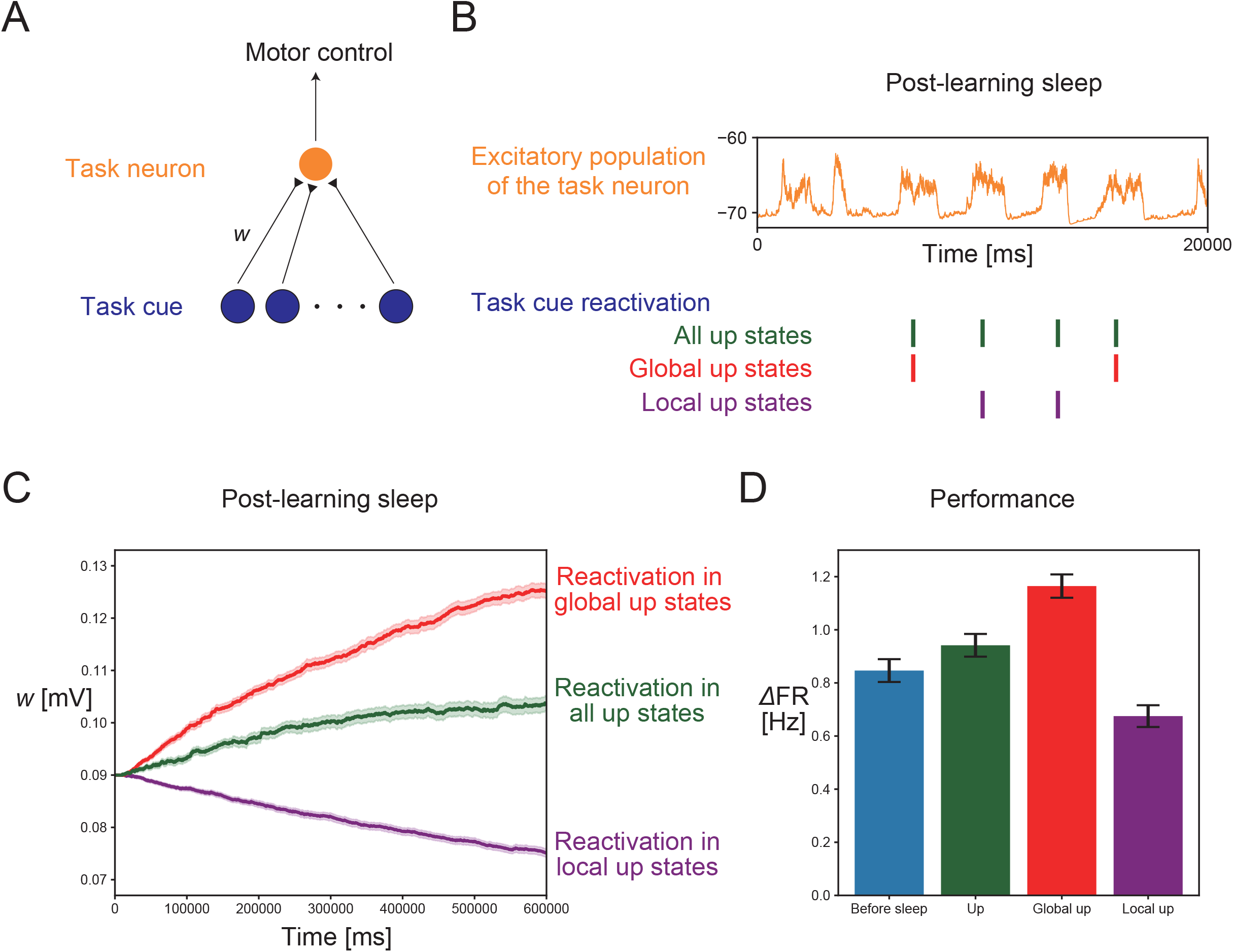
Changes in synaptic weights and task performance during post-learning NREM sleep. (A) The schematic description of the neuronal networks related to the task. Presynaptic neurons emitted synchronous spikes (“task cue”) during the task and also during the up states of the post-learning NREM sleep. The postsynaptic neuron (“task neuron”) is an excitatory neuron in the E1 population that is projected by the presynaptic neurons. Task performance is defined as the firing rate increase of the task neuron during the task period. (B) A typical mean membrane potential of the E1 population. In different simulations, the presynaptic neurons were stimulated during all the up states, global up states, or local up states of NREM sleep by neuronal reactivation. (C) The synaptic changes during the post-learning sleep. The synapses were potentiated by reactivation during the global up states or during all up states, whereas they were depressed by reactivation during the local up states. The lines and shadows represent mean and SEM, respectively. (D) The comparison of task performance before and after synaptic changes during sleep. This tendency is the same as that of the synaptic weight changes shown in Fig. 5C. Error bars represent SEM.

The feedforward synaptic weights were further potentiated in the simulation of post-learning NREM sleep (Fig. 5C). Task performance increased during sleep, reflecting this synaptic potentiation, consistent with the experimentally suggested memory consolidation during NREM sleep (Fig. 5D). Next, to investigate the roles of global and local slow waves in memory reorganization separately, we ran simulations where memory reactivation was restricted during either the global or local up states (Fig. 5B). The synaptic potentiation was enhanced when reactivation was restricted to global up states. Further, synaptic depression was observed instead when reactivation was restricted in local up states (Fig. 5C). As a result, the task performance showed a greater increase in the former case but a decrease in the latter case (Fig. 5D). The results were consistent with experimental findings that global and local slow waves contribute to memory consolidation and forgetting, respectively. These results suggest that the balance of global and local slow waves with distinct information transfer capacities regulates the spectrum of memory consolidation and forgetting via the infomax synaptic plasticity rule.

## 3 Discussion

Using a top-down approach with the information theory, we provided a unified learning rule, the infomax rule, for the state-dependent synaptic plasticity during NREM sleep. The infomax rule is comprised of the synaptic changes by the information term and synaptic depression by the synaptic cost term (Fig. 6A). A high firing rate condition biases the synaptic plasticity toward depression. The reason is that the signal-to-noise ratio for the synaptic transmission declines with the background firing rate of the postsynaptic neuron, and the cost term dominates the information term under a high firing rate condition. The learning rule yields an information-theoretical interpretation of different STDP observed in the up and down states [20]. Moreover, it also provides the distinct STDP during the global and local slow waves, suggesting a possible mechanism for balancing memory consolidation and forgetting. These properties are consistent with the role of neuronal reactivation in global and local slow waves [13].

**Figure 6.**
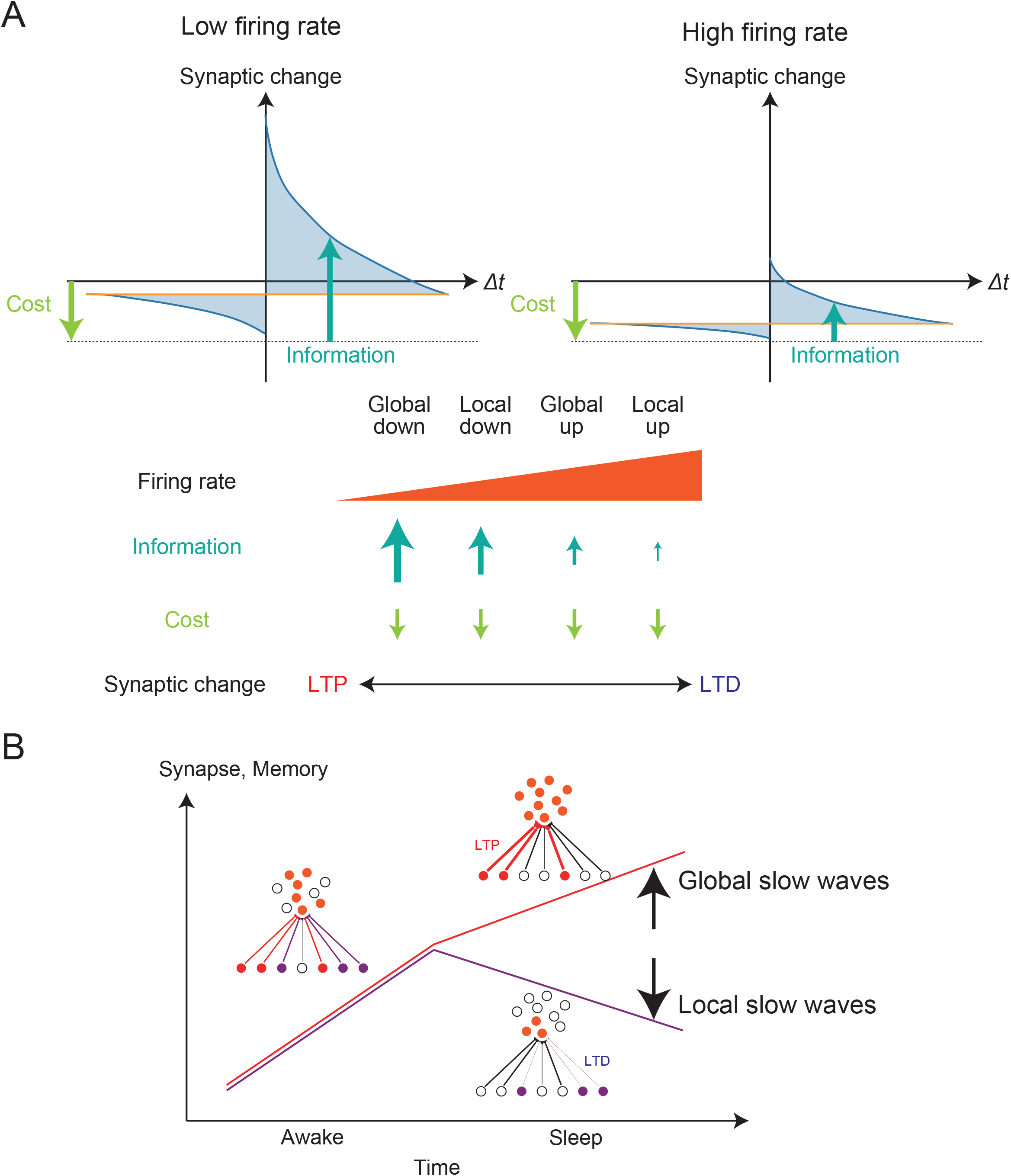
The proposed role of NREM sleep is to bridge neural information coding, synaptic plasticity, and memory reorganization. (A) The relationship between the mean firing rate and synaptic changes by the infomax rule. Synaptic potentiation by the information term was decreased at a high firing rate owing to many background spikes, whereas synaptic depression by the cost term was unaffected. Therefore, high firing rate induced the synaptic depression. Because the mean firing rates are global down < local down < global up < local up in ascending order, the amount of synaptic changes follows the opposite order. (B) The possible distinct roles of global and local slow waves. Reactivated patterns during global slow waves induced synaptic potentiation, whereas those during local slow waves induced synaptic depression. This could cause selective memory consolidation and forgetting of the reactivated patterns during the global and local slow waves, respectively.

The infomax rule not only reproduces the biased STDP toward depression during up states [20] but also provides a reliable prediction of the entire STDP curve during up states; the original experiment measured the synaptic change using a few representative time differences (10, 50, and −10 ms) between the presynaptic and postsynaptic spikes. The infomax rule further predicts that the STDP curve is sensitive to the initial synaptic weights and number of synchronous inputs (Fig. S2). These predictions are experimentally testable using protocols similar to those of [20], using various time differences, and altering the strength of presynaptic stimulations. Importantly, the infomax rule also predicts that relatively weak synapses can be potentiated, even in the up state, if a postsynaptic spike immediately follows a presynaptic spike (Fig. S2).

The infomax rule requires estimating the expected firing rate of each postsynaptic neuron in real-time to set the activity threshold separating the synaptic potentiation and depression. There are several biologically plausible implementations for computing this. The simplest estimate uses a temporally averaged firing rate. However, it tends to lag behind the true instantaneous firing rate in the presence of slow waves. In our slow-wave model, the expected instantaneous firing rate was accurately estimated using the average excitatory firing rate of the local network population. It is possible that the inhibitory neurons projected by nearby excitatory neurons compute the average firing rate, which is consistent with the observation that inhibitory input could modulate the balance of synaptic potentiation and depression [35]. Alternatively, astrocytes may temporally and spatially integrate nearby synaptic inputs and regulate the activity threshold separating synaptic potentiation and depression [36].

The infomax rule suggests the potential importance of down states for memory consolidation. Although some studies have assumed down states as resting periods for cellular maintenance [37], it has been reported that the hyperpolarization might be essential for slow waves to induce synaptic potentiation [38]. A recent study further suggested that the neuronal activities during down states (“delta spikes”) coincided with the hippocampal ripple activities and might be important for memory consolidation [34]. The infomax rule suggests that down states with fewer background spikes promote synaptic potentiation more than up states, implying that the delta spikes could effectively induce synaptic potentiation and memory consolidation. Many studies have also focused on the involvement of neuronal activities in up states or the transition period from down to up states during memory consolidation [39, 40]. Such distinct roles of the neuronal reactivation during the up and down states warrant further study.

The infomax rule in our sleep model suggests the importance of the spatial scale of slow waves [12, 41]. Specifically, the present model suggests that the synaptic changes induced by global slow waves are dominated by potentiation as compared to local slow waves because of different baseline firing rates. These predictions provide a unified view of apparently contradictory hypotheses regarding synaptic plasticity during sleep. The sleep synaptic homeostasis hypothesis [3] states that synaptic strength is down-scaled during sleep, which may promote memory consolidation by increasing the signal-to-noise ratio. On the other hand, other studies have reported synaptic potentiation and the formation of new spines during sleep [42, 43], which might be related to slow waves [38, 44]. The infomax rule can naturally induce the former effect during local slow waves and the latter during global slow waves, based on the information-theoretical principle.

Our theory could be verified experimentally along with the following two points. First, excitatory firing rates during global up states were lower than those during local up states because the surrounding neuronal activities increased the local inhibitory activity more greatly than the local excitatory activity in our model. Although the direct evidence supporting this prediction has not been reported to our knowledge, it could be indirectly supported by theoretical studies outside the sleep research. The dynamics of up states in this model are characterized as ISNs [32, 33, 31, 45], in which the unstable recurrent excitatory activities are stabilized by the local inhibitory circuits. To explain the surround suppression observed in the visual cortex, ISN models assume stronger excitatory inputs from the surrounding populations to local inhibitory populations than to local excitatory populations [33, 31]. Our model inherits this assumption. Furthermore, recent technical advances in neuronal recordings from a large number of neurons [46] can annotate the up or down states in each local area more precisely. In the future, such recordings could directly verify the model prediction that the mean firing rates of excitatory neurons decline with the spatial scale of the slow waves. Another testable model prediction is that global slow waves should permit more efficient information transmission than local slow waves. For example, one could optogenetically stimulate a small group of neurons and quantify the accuracy of stimulus encoding in neurons postsynaptic to the stimulated neurons during global and local slow waves.

The suggested role of memory consolidation during global slow waves and forgetting during local slow waves raises the possibility that there are mechanisms regulating the spatial scales of slow waves for selecting a subset of reactivation events to be consolidated (Fig. 6B). One possibility is that some neuronal populations projecting broadly to cortical neurons promote global synchronization. Previous studies have consistently suggested that the thalamus [47] and claustrum [48] play a role in synchronizing the down states of multiple cortical neurons. If such neuronal populations are co-active with reactivation patterns, these patterns could be selectively consolidated. Considering the high temporal correlation between the hippocampal sharp-wave ripples (SWRs) and cortical slow waves [49], specific neuronal populations may regulate both the generation of slow waves and neuronal reactivation. This possibility needs to be explored further.

In summary, the proposed theory bridges neuronal information coding, synaptic plasticity, and memory reorganization. Our normative framework provides a versatile learning rule for state-dependent synaptic plasticity and memory reorganization during NREM sleep.

## 4 Material and Methods

### 4.1 Spiking neuron model

We introduced a stochastic spiking neuron model; each neuron was either excitatory (E) or inhibitory (I). The spikes of each neuron in population P (P = E, I) were generated probabilistically with density *ρ*(*t*), as follows:

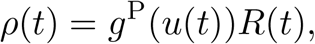

where *g*^P^(*u*) denoted a softplus activation intensity function, *u*(*t*) the membrane potential, and *R*(*t*) a refractory factor representing transient suppression of the instantaneous postsynaptic firing rate after a postsynaptic spike. The function 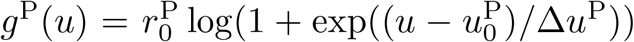 with 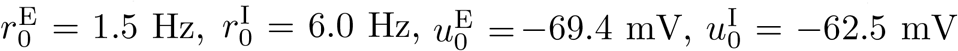, and Δ*u*^E^ = Δ*u*^I^ = 0.5 mV. The refractory factor is modeled to be the same for excitatory and inhibitory neurons as 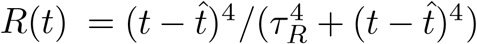, where 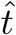 denoted the last spike time of the postsynaptic neuron and time constant *τ*_*R*_ = 30 ms. The suppression of this factor after spiking may reflect several mechanisms, including classical refractoriness [50], afterhyperpolarization (AHP) [51], and EPSP suppression by back-propagating action potential [52].

### 4.2 Feedforward model

We modeled a postsynaptic neuron that received the feedforward inputs from *N* presynaptic excitatory neurons. Each spike of presynaptic neuron *j* evoked EPSP of amplitude *w*_*j*_ and exponential time course 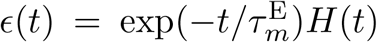 with time constant 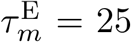 ms and Heaviside step function *H*(*t*). The EPSP time course from presynaptic neuron *j* was denoted by 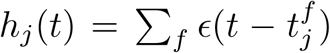, summing the influence of presynaptic spikes at time 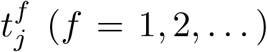. Then, the postsynaptic membrane potential *u*(*t*) was denoted by

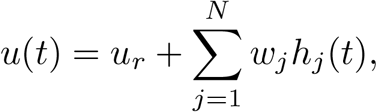

with resting membrane potential *u*_*r*_ = −70 mV. Postsynaptic spikes were generated probabilistically with density *ρ*(*t*) = *g*^E^(*u*(*t*))*R*(*t*), as mentioned in the previous section.

### 4.3 Information maximizing learning rule

We used the infomax rule for synaptic plasticity [24, 25]. The objective function *L* was described as:

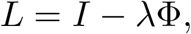

with the mutual information term *I*, cost term Φ of the synaptic weights, and coefficient parameter *λ* = 0.32 [1*/*(mV)^2^]. *I* measures the mutual information between the pre- and postsynaptic spike trains. We omitted the homeostatic term included in previous studies because it did not contribute to our results. Each term was denoted by

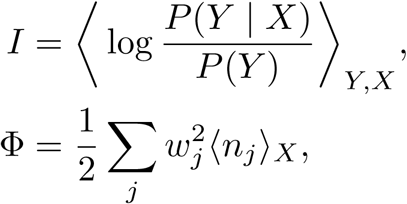

with presynaptic and postsynaptic spike trains *X* and *Y*, respectively, and *n*_*j*_ represents the number of presynaptic spikes at synapse *j* during duration *T*. The presynaptic and postsynaptic spike trains up to time *t* were formally denoted by 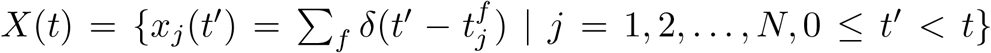 and 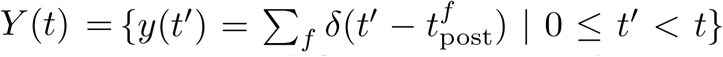, respectively, where 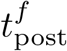 represents the *f*-th (*f* = 1, 2, …) postsynaptic spike timings. We specifically wrote *X* = *X*(*T*) and *Y* = *Y* (*T*) to represent the entire spike train from time *t* = 0 to *t* = *T*. Note that the angular brackets ⟨·⟩_*Y,X*_ and ⟨·⟩_*X*_ represent the averages over all possible *Y, X*, and *X*, respectively.

The optimal synaptic weight change followed the gradient ascent algorithm

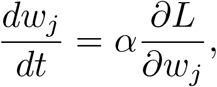

with a learning rate *α* = 0.01(mV)^2^. By calculating the gradient [24, 25], the infomax rule was described as

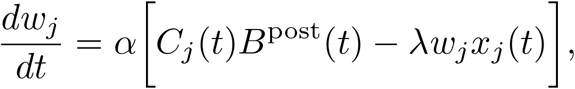

with

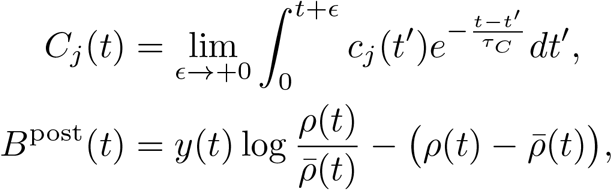

described with auxiliary variable 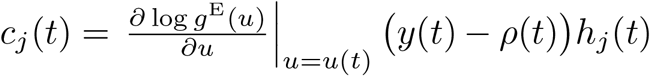, expected firing rate 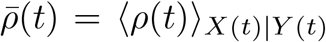 and time constant *τ*_*C*_ = 100 ms. The expected firing rate was further computed as

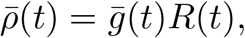

using the expected intensity 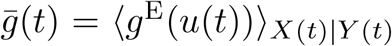; however, this value can be difficult to calculate if 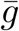 is time dependent. We estimated 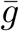 using slightly different methods for the feedforward and slow-wave models, as described in the corresponding sections.

### 4.4 Simulation of STDP in the feedforward model

We considered two populations of presynaptic neurons projecting onto a post-synaptic neuron. One population represented the stimulated neurons, while the other represented non-stimulated neurons. *N*_1_ = 100 non-stimulated neurons had fixed synaptic weight *w*_*j*_ = 0.5 mV and emitted spikes at 0.1 Hz in down state and 2.0 Hz in up states. *N*_2_ = 20 stimulated neurons had a plastic synaptic weight, which was initialized at *w*_*j*_ = 0.5 mV and updated by the infomax rule, and spiked only when they were stimulated. In Fig. S2, we varied the *N*_2_ value and the initial value of *w*_*j*_ in the stimulated neurons. STDP stimulation with time interval Δ*t* evoked spikes in all the stimulated neurons at time 0 and evoked a postsynaptic spike at time Δ*t*. The STDP stimulations were given twice 5000 ms apart. In this model, 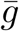 was estimated by averaging *g*(*u*(*t*)) over fluctuating *u*(*t*), while the stimulated presynaptic neurons remained silent and the non-stimulated presynaptic neurons spontaneously generated Poisson spikes at different firing rates for the up or down states.

### 4.5 Theoretical analysis of the STDP effect

For the theoretical analyses in this section, we approximated the softplus activation function using a linear function *g*^E^(*u*) = *g*_0_ · (*u* − *u*_*r*_). We evaluated the synaptic changes due to the pre-post stimulation with synchronous presynaptic spikes at *t* = 0 and a postsynaptic spike immediately afterward (at *t* = Δ*t >* 0 with the limit of Δ*t* → 0). To estimate the effect of STDP analytically and qualitatively, we made some simplifications. We approximated that the refractory factor *R* abruptly recovered from zero to one duration *τ*_*R*_ after the postsynaptic spike, and that the spontaneous spiking of the non-stimulated presynaptic neurons yielded a constant baseline membrane potential *u*_0_. We also assumed that, after presynaptic stimulation at *t* = 0, the membrane potential *u*(*t*) = *u*_0_ + Σ_*j∈*stim_ *w*_*j*_*h*_*j*_(*t*) decayed back to the baseline *u*_0_ by the time the postsynaptic neuron recovered from the refractoriness because 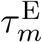 was smaller than *τ*_*R*_. Here, stim denotes the set of stimulated neurons. The baseline intensity is denoted by 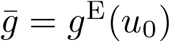, and the peak intensity increase after presynaptic stimulation by 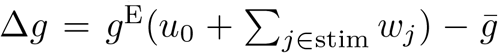. In this case, we found 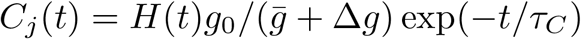 and 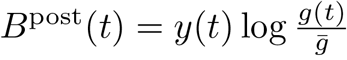. Note that no extra postsynaptic spike was possible when *h*_*j*_ was significantly positive because the refractory period was longer than that of the stimulus-caused EPSP duration. Hence, the change in the *j*-th synaptic weight due to the STDP stimulus was

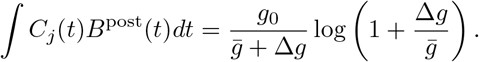

This result indicated that the increase in synaptic weights due to the information term decreased with the mean firing intensity 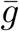.

### 4.6 Slow wave model

We considered *N* ^E^ = 800 excitatory neurons, E, and *N* ^I^ = 200 inhibitory neurons, I, divided into four local networks. Each local network contained 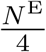 excitatory neurons and 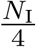 inhibitory neurons. Within each local network, no connections exist between two inhibitory neurons, and all-to-all connections exist between two excitatory neurons as well as between excitatory and inhibitory neurons. Between different local networks, the connection probability from E to E was *p*_EE_ and the connection probability from E to I was *p*_IE_, whereas inhibitory neurons did not send long-range connections to the other local networks. The connection probabilities *p*_EE_ and *p*_IE_ were set to 0.05 and 0.3, respectively, except for those in Figs. 3C and S1A. There is no self-coupling in these neurons. The dynamics of the membrane potential 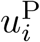 of neuron *i* in population P (P = E, I) were described by

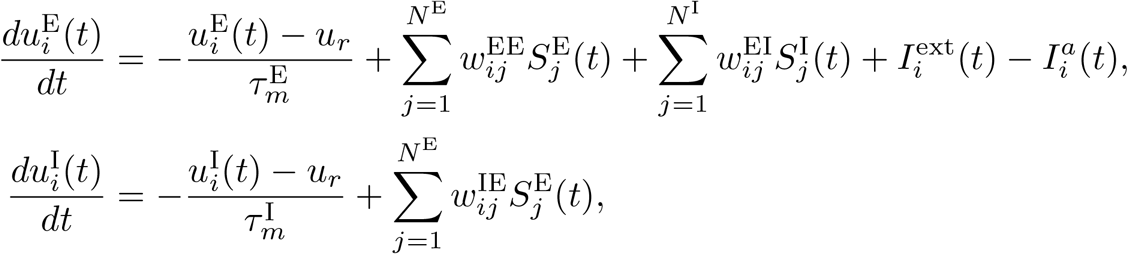

with the recurrent synaptic weight 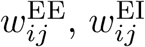, and 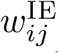 from excitatory neuron *j* to excitatory neuron *i*, from inhibitory neuron *j* to excitatory neuron *i*, and from excitatory neuron *j* to inhibitory neuron *i*, respectively, membrane time constant 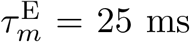 for excitatory neurons and 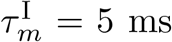 for inhibitory neurons, the resting membrane potential *u*_*r*_ = −70 mV, spike train 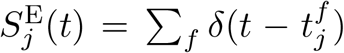 or 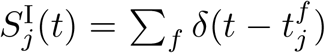 of excitatory or inhibitory neuron *j* described with its *f* -th spike time 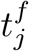, external current 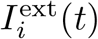, and adaptation current 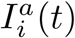. The spikes of neuron *i* in population P (P = E, I) were generated probabilistically with an instantaneous firing rate 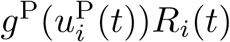, as mentioned in the previous section, with its refractory factor *R*_*i*_. The recurrent synaptic weight was fixed at 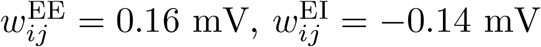, and 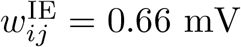 if synaptic connections existed from neuron *j* to neuron *i*, whereas it was set to 0 mV if there were no synaptic connections. The external current was a feedforward input only to the excitatory neuron 1, described as 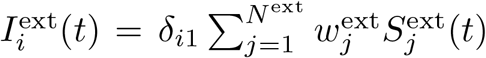, with the Kronecker delta *δ*_*i*1_, synaptic weights 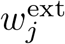 from *N* ^ext^ presynaptic neurons, and presynaptic spike train 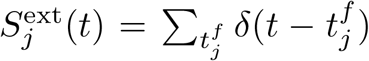, where 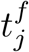 represented the spike timing. Presynaptic neurons emitted the synchronous spikes upon STDP stimulation (Fig. 4) or task stimulation (Fig. 5), although they did not emit spontaneous spikes. The STDP and task simulations (Figs. 4 and 5) changed the feedforward synaptic weights *w*_*j*_ according to the infomax rule. The dynamics of the adaptation current 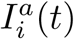 of neuron *i* were described as follows:

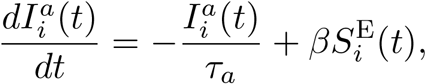

with a time constant of *τ*_*a*_ = 1500 ms and a constant value of *β* = 0.0077 mV/ms.

### 4.7 stage classification

First, we classified the up and down states of each population using two transition thresholds. When the mean membrane potential of excitatory neurons in each local network exceeded the up-transition threshold *θ*_up_ = −68.25 mV, this moment was judged as a state transition to the up state. When the mean membrane potential fell below the down-transition threshold *θ*_down_ = −69.75 mV, this moment was judged as a state transition to the down state. We then classified each state into the global or local states. When a local network was in the down state, it was classified into the global down state if the number of other local networks in the down states was two or three, while it was classified as the local down state otherwise. Likewise, when a focused population was in the up state, it was classified into the global up state if the number of other local networks in the up states was two or three, while it was classified as the local up state otherwise.

### 4.8 Phase plane analysis

Assuming a large population of neurons in each local network, the average dynamics of the membrane potential *u*^P^ (P = E, I) in the population P (containing 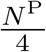 neurons) was described by

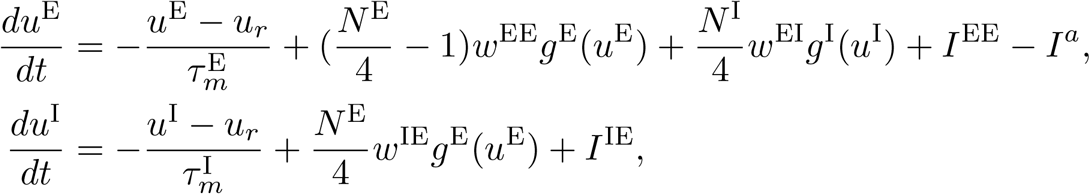

with the synaptic weight *w*^EE^ = 0.16 mV, *w*^EI^ = −0.14 mV, *w*^IE^ = 0.66 mV, and *I*^PE^ denoting the long-range excitatory currents to population P from the other local networks. External inputs were not considered here. We denoted by 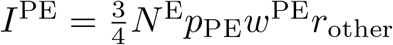 the excitatory input from the other local networks to population P, where *r*_other_ represented the firing rate of the excitatory populations in the other local networks. *r*_other_ was set to 0 Hz and 6 Hz when the other local networks were in the down and up states, respectively. The adaptation current *I*^*a*^ was set to 0, 0.01, 0.05, or 0.18 mV/ms.

Furthermore, we conducted a stability analysis in the slow wave model. We approximated the activation function *g*^P^(*u*) (P = E, I) using the threshold-linear function 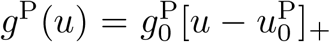, where 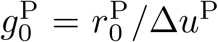 and [*x*]_+_ took the value *x* if *x >* 0 and zero otherwise. (Note that the phase plane plots in Figs. 3G and Fig. S1B are depicted using the original softplus activation function.) By using this approximation, the differential equations above were described as

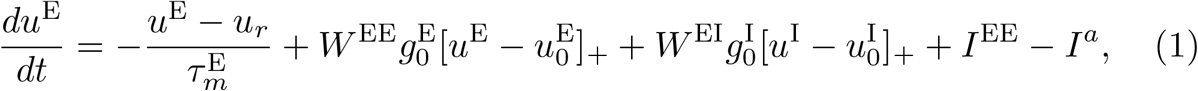

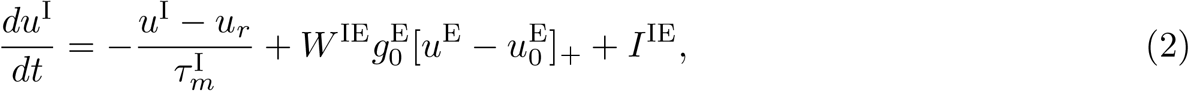

where 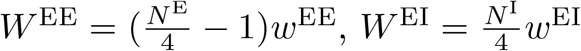, and 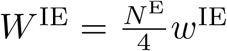.

We first investigated the case in which the external currents *I*^EE^ and *I*^IE^ were zero. We searched the condition that both the down state satisfying 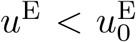 and 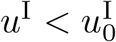 and the up state satisfying 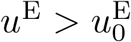 and 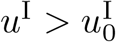 existed stably.

If a solution satisfied 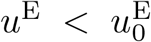 and 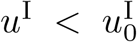, then the fixed point was 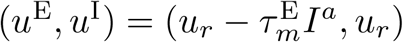. Hence, this solution existed when it satisfied

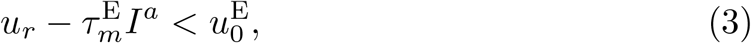

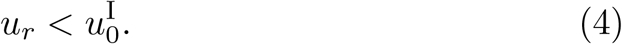

The Jacobian matrix *J* of differential equations (1) and (2) at the fixed point satisfying 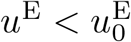 and 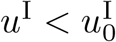 was described as

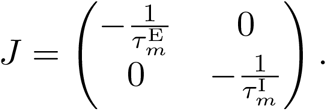

Because all eigenvalues of the Jacobian matrix *J* had a negative real part, the fixed point was stable.

If a solution satisfied 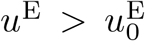 and 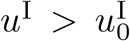, then the fixed point of the equations was

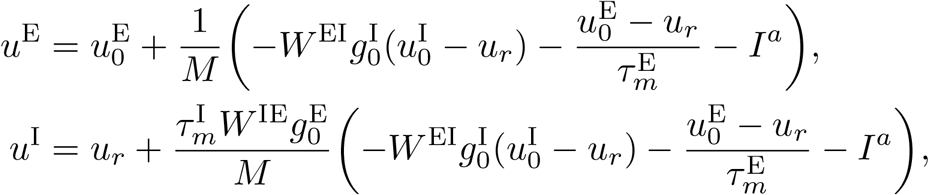

where 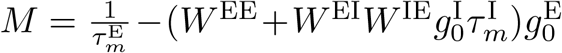. This solution existed when 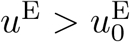 and 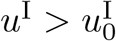 were satisfied. Under the condition (4), it was equivalent to

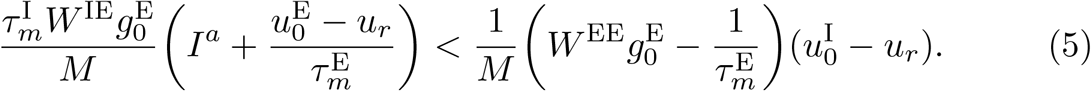

The Jacobian matrix *J* of differential equations (1) and (2) at the fixed point satisfying 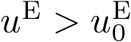 and 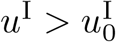 was described as

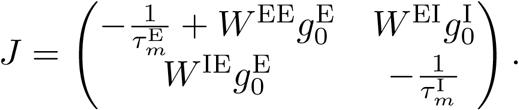

If all eigenvalues of the Jacobian matrix *J* had a negative real part, then the fixed point was stable. This condition was equivalent to having a negative trace and a positive determinant of matrix *J*; namely,

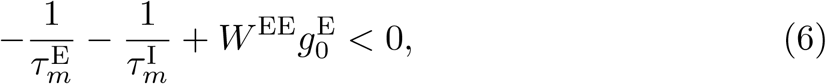

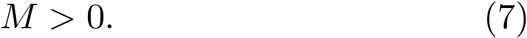

In summary, two stable states existed when the conditions (3)–(7) were satisfied.

Next, we considered the case where the external currents *I*^EE^ and *I*^IE^ were not zero. We investigated the condition in which the long-range excitatory input from the other local networks suppressed the firing rates of the local excitatory population in the up state. The fixed point corresponding to the up state was

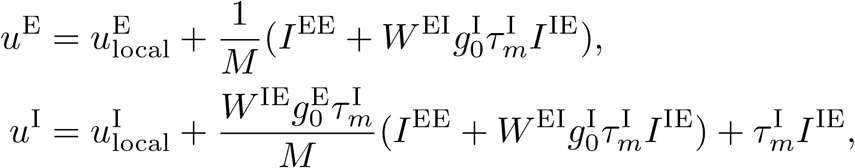

where 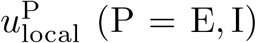 was a fixed point when external currents *I*^EE^ and *I*^IE^ were zero. Hence, the condition in which the external currents suppressed the excitatory firing rates was described as:

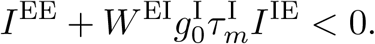

Using 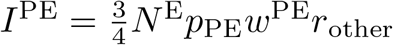, the condition was further transformed into:

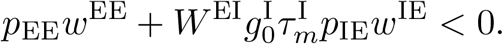

### 4.9 Simulation of STDP in the slow wave model

In the slow-wave model, the expected firing intensity 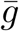(*t*) was estimated by the population-mean firing intensity of the excitatory neurons in the same population, described as

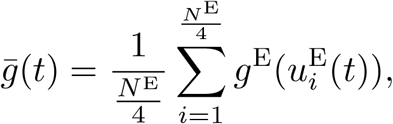

where the excitatory neurons 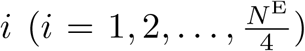 were in the same excitatory population (E1) as the postsynaptic neuron in consideration. In Fig. 4, we considered *N* ^ext^ = 20 presynaptic neurons that had the plastic synaptic weights with an initial value of 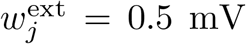. The presynaptic neurons emitted no spontaneous spikes. We considered STDP stimulations with a time difference of −80 ≤ Δ*t* ≤ 80 ms and pre-only stimulations when the E1 population was in each state (global up, local up, global down, or local down state). In the case of the pre-post stimulation (i.e., Δ*t* is positive), the stimulation evoked synchronous presynaptic spikes first followed by Δ*t*, a postsynaptic spike. Stimulation was applied if the target state of the E1 population continued to be more than 200 ms, and the interval from the last stimulation was 500 ms or more at the candidate time of the presynaptic stimulation. In the case of the post-pre-stimulations (i.e., Δ*t* is negative), a postsynaptic spike was evoked Δ*t* before the synchronous presynaptic spikes. Stimulation was applied if the above condition was satisfied at the candidate time of the postsynaptic stimulation. The pre-only stimulation was the same as the pre-post stimulation, except that the postsynaptic stimulation was absent. The number of stimulations was fixed at ten. To exclude rare network configurations due to random synaptic connections in which the target states hardly appear, the simulation time was fixed to 50000 ms, and the simulations in which the number of stimulations did not reach ten by the end of the simulation time were excluded from the analysis.

### 4.10 Task simulation

Fig. 5 described the task simulation. The model connection structure was similar to that shown in Fig. 4, containing *N* ^ext^ = 40 presynaptic neurons that had plastic synaptic weights with an initial value of 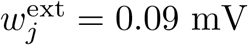 (Fig. 5A).

First, we introduced the awake period. While awake, the adaptation current was set to be constant at 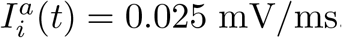, in which the neurons showed a continuing depolarized membrane potential similar to the awake firing patterns experimentally observed. In the absence of task-related stimulation, the mean excitatory firing rate during wakefulness was approximately 6.0 Hz. We referred to this firing rate as the baseline firing rate.

Next, we considered a task inspired by the brain-machine-interface task [13] while awake. We assumed that the presynaptic neurons emit synchronous Poisson spikes at 5 Hz during the task. Task performance was defined as the difference between the mean firing rate of the postsynaptic neuron during the task period and the baseline firing rate. In this setting, the feedforward synaptic weights contributed to the task performance.

Finally, we considered the synaptic changes and improvements in task performance during the post-learning NREM sleep period. During NREM sleep, the dynamics of the adaptation current followed the slow-wave model in the previous section, and the presynaptic neurons emitted synchronous Poisson spikes at 5 Hz as a neuronal reactivation during the global and/or local up states of the E1 population. The synaptic weight changed according to the infomax rule. The task performance was quantified by the above measure using the synaptic weights before and after NREM sleep.

### 4.11 Simulation environment

All numerical calculations were performed using the custom-written Python codes. The model was simulated in discrete time with time steps of 1 ms. Synaptic weights did not change during the first 10000 ms in all simulations to avoid the effects of initial values. The initial value of the last spike time was set to −10000 ms for all neurons. The source code will be appended as Supplementary Information upon publication.

## 5 Figure legends

**Figure S1.**
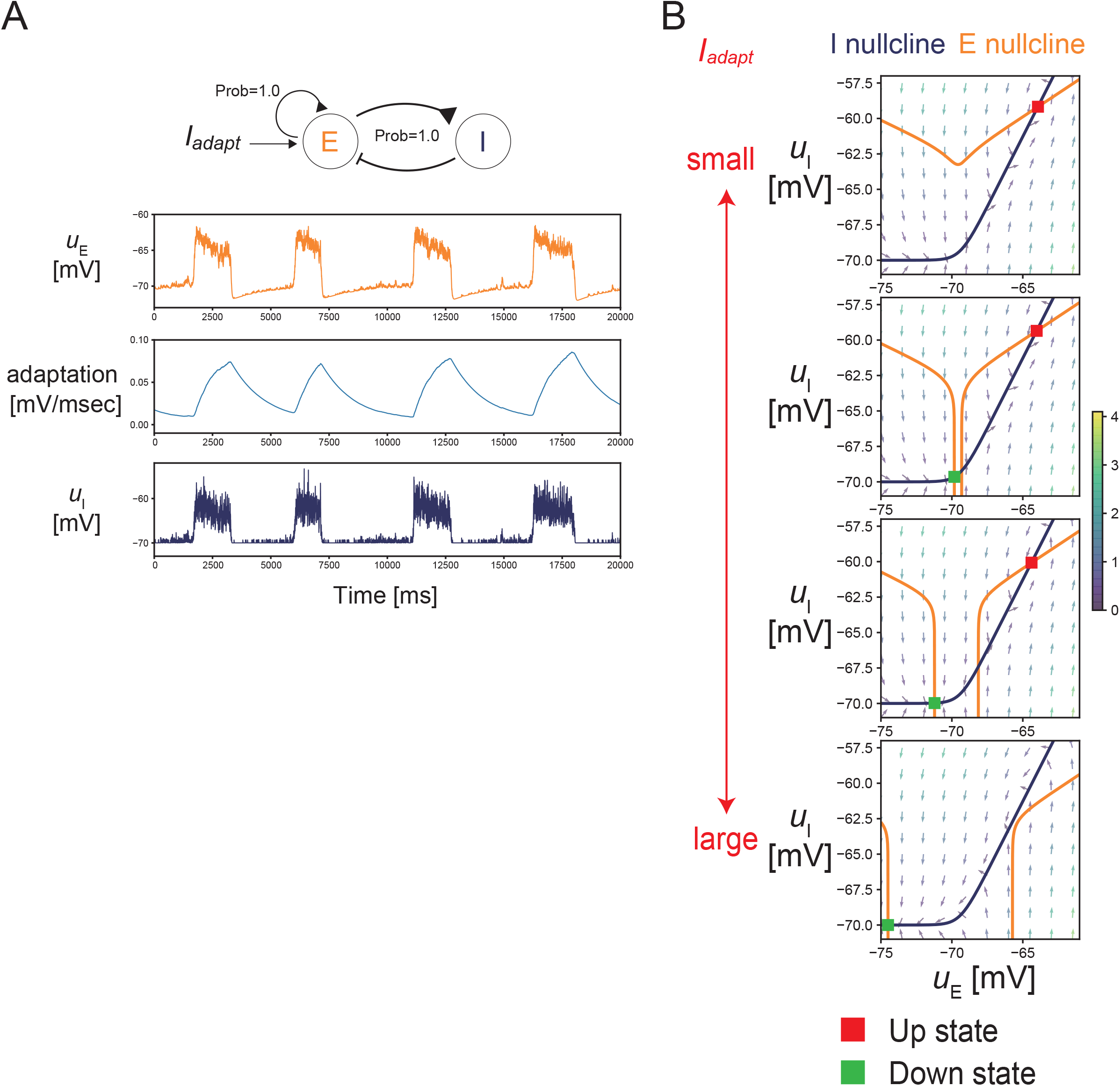
The spatially homogeneous model with 200 excitatory and 50 inhibitory neurons. (A) The membrane potential of excitatory and inhibitory neurons transit between up and down states with changing adaptation currents. (B) The phase plane plot of the model. The excitatory nullcline changed with distinct adaptation currents.

**Figure S2.**
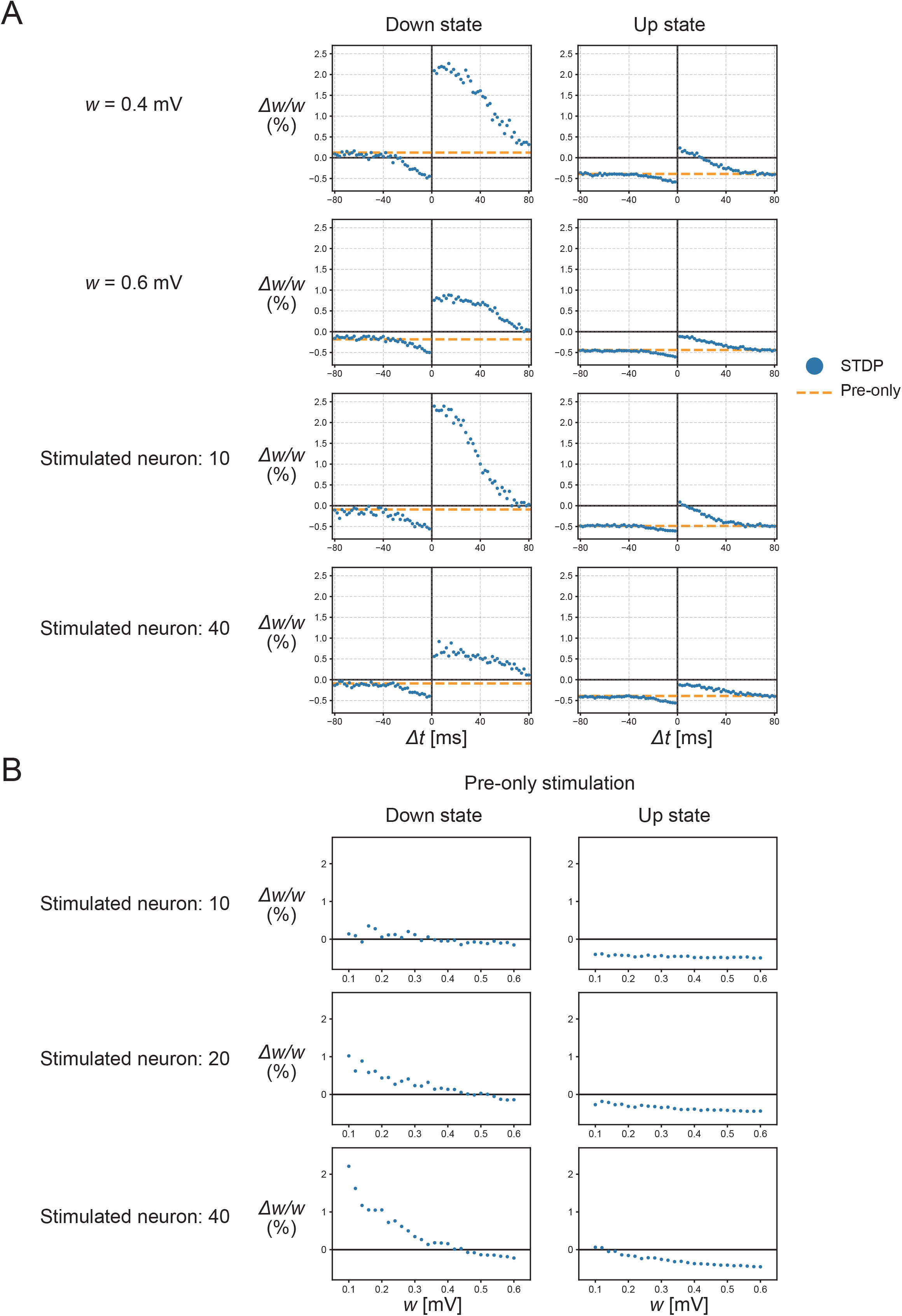
STDP curves for distinct initial synaptic weights and numbers of stimulated neurons in the feedforward model. (A) The STDP curves in distinct synaptic weights and the distinct numbers of the stimulated neurons. (B) The synaptic weight changes by the pre-only stimulations. The smaller initial synaptic weights and larger number of stimulated neurons bias the changes toward potentiation.

## 6 Acknowledgments

The authors thank Hideaki Kume, Shoi Shi, and Genki Shimizu for their helpful discussion. This study was supported by RIKEN Center for Brain Science, Brain/MINDS from AMED under Grant No.JP15dm0207001 (T.T.), KAKENHI Grant-in-Aid JP18H05432 (T.T.) and JP21J10564 (K.Y.) from JSPS, RIKEN Junior Research Associate Program (K.Y.), and Masason Foundation (K.Y.).

## 7 Author contributions

K.Y. and T.T. conceived the project. K.Y. conducted numerical simulations. K.Y. and T.T. wrote the manuscript.

## 8 Data availability

The source code will be appended as Supplementary Information upon publication.

